# Reinforcement Learning approaches to hippocampus-dependent flexible spatial navigation

**DOI:** 10.1101/2020.07.30.229005

**Authors:** Charline Tessereau, Reuben O’Dea, Stephen Coombes, Tobias Bast

## Abstract

Humans and non-human animals show great flexibility in spatial navigation, including the ability to return to specific locations based on as few as one single experience. To study spatial navigation in the laboratory, watermaze tasks, in which rats have to find a hidden platform in a pool of cloudy water surrounded by spatial cues, have long been used. Analogous tasks have been developed for human participants using virtual environments. Spatial learning in the watermaze is facilitated by the hippocampus. In particular, rapid, one-trial, allocentric place learning, as measured in the Delayed-Matching-to-Place (DMP) variant of the watermaze task, which requires rodents to learn repeatedly new locations in a familiar environment, is hippocampal dependent. In this article, we review some computational principles, embedded within a Reinforcement Learning (RL) framework, that utilise hippocampal spatial representations for navigation in watermaze tasks. We consider which key elements underlie their efficacy, and discuss their limitations in accounting for hippocampus-dependent navigation, both in terms of behavioural performance (*i.e*., how well do they reproduce behavioural measures of rapid place learning) and neurobiological realism (i.e., how well do they map to neurobiological substrates involved in rapid place learning). We discuss how an actor-critic architecture, enabling simultaneous assessment of the value of the current location and of the optimal direction to follow, can reproduce one-trial place learning performance as shown on watermaze and virtual DMP tasks by rats and humans, respectively, if complemented with map-like place representations. The contribution of actor-critic mechanisms to DMP performance is consistent with neurobiological findings implicating the striatum and hippocampo-striatal interaction in DMP performance, given that the striatum has been associated with actor-critic mechanisms. Moreover, we illustrate that hierarchical computations embedded within an actor-critic architecture may help to account for aspects of flexible spatial navigation. The hierarchical RL approach separates trajectory control via a temporal-difference error from goal selection via a goal prediction error and may account for flexible, trial-specific, navigation to familiar goal locations, as required in some arm-maze place memory tasks, although it does not capture one-trial learning of new goal locations, as observed in open field, including watermaze and virtual, DMP tasks. Future models of one-shot learning of new goal locations, as observed on DMP tasks, should incorporate hippocampal plasticity mechanisms that integrate new goal information with allocentric place representation, as such mechanisms are supported by substantial empirical evidence.

## 1 Introduction

Successful spatial navigation is required for many everyday tasks: animals have to find food and shelter and remember where and how to find these, humans need to navigate to work, home or to the supermarket. Since natural environments are inherently heterogeneous and subject to continuous change, animal brains have evolved robust and flexible solutions to solve challenges in spatial navigation.

In the mammalian brain, and to some extent also the avian brain (Colombo and Broadbent 2000;Bingman and Sharp 2006), the hippocampus has long been recognised to play a central role in place learning, memory and spatial navigation, based on the behavioural effects of lesions and other manipulations of the hippocampus (Morris et al. 1982, 1986, 1990) and based on the spatial tuning of certain hippocampal neurons, so-called place cells (O’Keefe and Dostrovsky 1971; Moser et al. 2017; O’Keefe 2014; Jeffery 2018). Studies combining hippocampal manipulations with behavioural testing in rodents have revealed that the hippocampus is particularly important for flexible spatial navigation based on rapid allocentric place learning, in which places are learned based on their relationship to environmental cues (Morris et al. 1990; Eichenbaum 1990; Steele and Morris 1999; Bast et al. 2009).

Animal experiments on spatial navigation have been complemented by tools from theoretical and computational neuroscience. Many theoreticians have targeted spatial navigation problems, either trying to reproduce behaviours (Dayan 1991;Banino et al. 2018) or to explain properties of neurons that show spatial tuning, including hippocampal place cells (Samsonovich and McNaughton 1997; O’Keefe and Burgess 1996; Fuhs and Touretzky 2006; Widloski and Fiete 2014; Banino et al. 2018). Some have also studied the functional properties of network representations of neural computations. For example, exploring how static attributes such as place coding can lead to or be integrated within network dynamics (Kanitscheider and Fiete 2017), or have probed the storage capacity of spatial representations (Battaglia and Treves 1998).

In a spatial navigation context, most real-world situations involve choosing a behavioural response that leads to a goal location associated with a reward. These could be direct rewards, such as food, or escape from an unpleasant situation, *e.g*. escape from water in the watermaze. Successful navigation requires animals to distinguish diverse cues in their environment, to encode their own current position, to access a memory of where the goal is, to choose an appropriate trajectory and to recognise the goal and its vicinity. Learning how to reach a goal location is a problem that, in principle, fits very well within a Reinforcement Learning (RL) context. RL commonly refers to a computational framework that studies how intelligent systems learn to associate situations with actions in order to maximise the rewards within an environment (Sutton and Barto 2018). When applied to spatial navigation, a RL model can be used to infer how neuronal representations of space, as revealed by electrophysiological recordings, may serve to maximise reward (Foster et al. 2000; Banino et al. 2018; Gerstner and Abbott 1997; Corneil and Gerstner 2015; Russek et al. 2017; Dollé et al. 2018). RL models have led to numerous successes in understanding how biological networks could produce observed behaviours (Haferlach et al. 2007; Frankenhuis et al. 2019), yet there are still substantial challenges in using RL approaches to account for animal behaviour, such as hippocampus-dependent flexible navigation based on rapid place learning.

The aim of this paper is to review RL models that may account for hippocampus-dependent rapid place learning, especially as seen in the watermaze DMP task. We will especially focus on an exemplar approach to this problem proposed by Foster et al. (2000). In section 2, we review briefly some key experimental findings on the involvement of the hippocampus in spatial navigation tasks in the watermaze, highlighting its particular importance in rapid place learning. Section 3 contains an overview of the key concepts in RL. Section 4 describes the first part of the model by Foster et al. (2000), a RL architecture that provides a computational approach to how a place can become associated with a reward. We present a detailed description of the computations underlying the behaviour of the model and their possible biological substrates, which we hope may make the model more accessible to neuroscientists without a strong neurocomputational background. Then, we focus on two minimal extensions to this architecture that enable adaptation to a changing reward location. The first one involves a map-like representation of location that enables vector-based navigation and was proposed by Foster et al. (2000), we show that this extension can reproduce key measures of rapid place learning performance on the DMP task, including sharp latency reductions from trial 1 to 2 (Steele and Morris 1999), and also the more recent finding that rats show search preference for the correct location within one trial (Bast et al. 2009). The second uses ideas drawn from hierarchical RL (Botvinick et al. 2009;Schweighofer and Doya 2003), in which adding layers of control allows the agent more flexible behaviours. We discuss details of these computations, their correspondence to neurobiological findings, and the plausibility of their implementation, in particular to account for rapid place learning within an artificial watermaze set-up. In the concluding section 5, we emphasise some of the computational principles that we propose hold particular promise for neuropsychologically realistic models of rapid place learning in the watermaze.

## 2 Flexible hippocampal spatial navigation

Humans and other animals show remarkable *flexibility* in spatial navigation. In this context, flexibility refers to the ability to adjust to a changing environment, such as the variation in the goal or start location (Tolman 1948). Watermaze tasks, in which rodents learn to find a hidden escape platform in a circular pool of water surrounded by spatial cues (Morris 2008), have been important tools to study the neuropsychological mechanisms of such flexible spatial navigation in rodents. In the original task, the platform location remains the same over many trials and days of training. The animals can incrementally learn the place of the hidden platform using distal cues surrounding the watermaze, and then navigate to it from different start positions (Morris 1981). Learning is reflected by a reduction in the time taken to reach the platform location (“escape latencies”) across trials and a search preference for the vicinity of the goal location when the platform is removed in probe trials.

Rapid place learning can be assessed in the watermaze through the delayed-matching-to-place (DMP) task, where the location of the platform remains constant during trials within a day (typically four trials per day), but is changed every day (Fig.1a, Steele and Morris (1999); Bast et al. (2009)). A key observation from the behaviour of rats on the DMP task is that a single trial to a new goal location is sufficient for the animal to learn this location and subsequently to navigate to it efficiently (Steele and Morris 1999). This phenomenon is therefore commonly referred to as “one-shot” or “one-trial” place learning. Such one-trial place learning is reflected by a marked latency reduction between the first and second trials to a new goal location (Fig.1b), with little further improvement on subsequent trials, and by a marked search preference for the vicinity of the correct location when trial 2 is run as probe (Fig.1c) with the platform removed (Bast et al. 2009).Buckley and Bast (2018) reverse-translated the watermaze DMP task into a task for human participants, using a virtual environment presented on a computer screen, and have shown that human participants exhibit similar one-trial place learning to rats.

**Figure 1.**
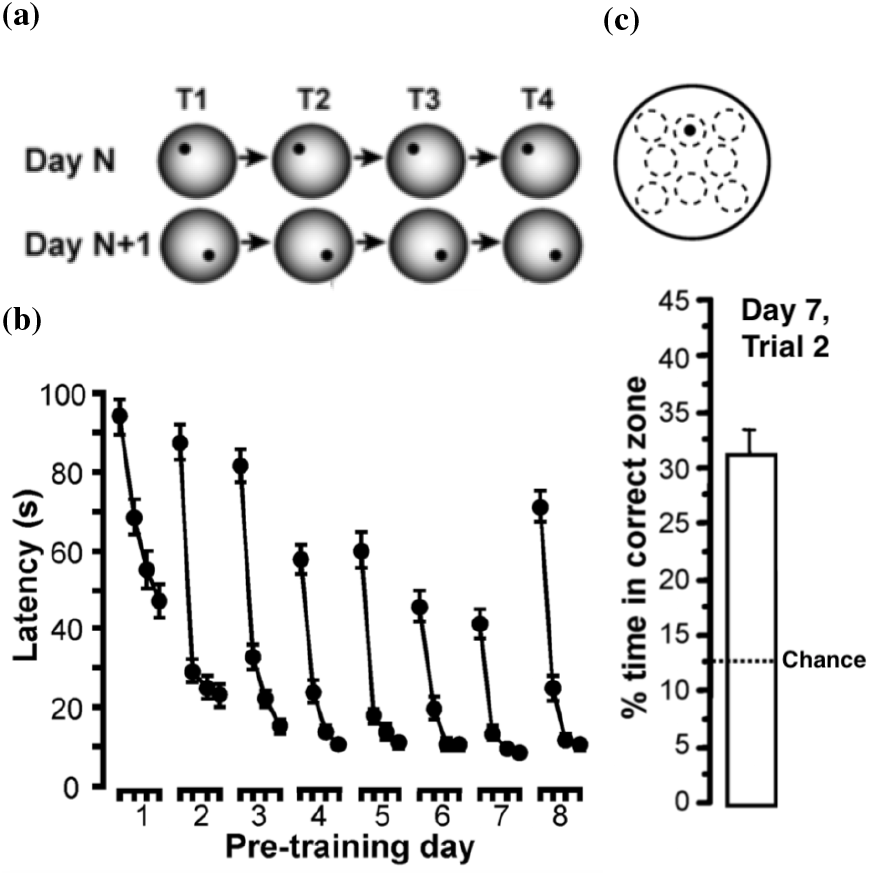
One-shot place learning by rats in the delayed-matching-to-place (DMP) watermaze task. (a) Rats have to learn a new goal location (location of escape platform) every day, and complete four navigation trials to the new location on each day. (b) The time taken to find the new location reduces markedly from trial 1 to 2, with little further improvements on trials 3 and 4, and minimal interference between days. (c) When trial 2 is run as a probe trial, during which the platform is unavailable, rats show marked search preference for the vicinity of the goal location. To measure search preference, the watermaze surface is divided in eight equivalent symmetrically arranged zones (stippled lines in sketch), including the ‘correct zone’ centred on the goal location (black dot). The search preference corresponds to time spent searching in the ‘correct zone’, expressed as a percentage of time spent in all eight zones together. The chance level corresponds to 12.5%, corresponding to the rat spending the same time in each of the eight zones depicted in the sketch. These behavioural measures highlight successful one-shot place learning. Figure adapted from Fig. 2 in (Bast et al.2009)

On the incremental place learning task in the watermaze, hippocampal lesions are known to disrupt rats’ performance (Morris et al. 1982), slowing down learning (Morris et al.1990) and severely limiting rats’ ability to navigate to the goal from variable start positions (Eichenbaum 1990). However, rats with partial hippocampal lesions sparing less than half of the hippocampus can show relatively intact performance on the incremental place learning task (de Hoz et al. 2003;Moser et al. 1995), and even rats with complete hippocampal lesions can show intact place memory following extended incremental training (Bast et al. 2009; Morris et al. 1990). Rats can also show intact incremental place learning on the watermaze with blockade of hippocampal synaptic plasticity if they received pretraining (Bannerman et al. 1995;Inglis et al. 2013). These findings suggest that incremental place learning, although normally facilitated by hippocampal mechanisms, can partly be sustained by extra-hippocampal mechanisms.

In contrast to incremental place learning, rapid place learning, based on one or a few experiences, may absolutely require the hippocampus, with extra-hippocampal mechanisms unable to sustain such learning (Bast 2007). Studies in rodents have shown that spatial navigation based on one-trial place learning on the DMP watermaze task is highly sensitive to hippocampal dysfunction that may leave incremental place learning performance in the watermaze relatively intact.

Specifically, one-trial place learning performance on the watermaze DMP test is severely impaired, and often virtually abolished, by complete and partial hippocampal lesions (Bast et al. 2009; De Hoz et al. 2005; Morris et al. 1990), as well as by disruption of hippocampal plasticity mechanisms (Inglis et al. (2013); Nakazawa et al. (2003); O’Carroll et al. (2006);Pezze and Bast (2012); Steele and Morris (1999), also compare similar findings by (Bast et al. 2005) in a dry-land food-reinforced DMP task) or by aberrant hippocampal firing patterns (McGarrity et al. 2017). Rats with hippocampal lesions and NMDA receptor blockade show similar swim paths on trial 1 and trial 2 to the same goal location, swimming in circles over large areas of the watermaze surface (Steele and Morris 1999; Redish and Touretzky 1998), suggesting that they do not have or cannot access information about the recent goal location and/or to the history of their positions.

Consistent with findings in rats that watermaze DMP performance is highly hippocampus-dependent, human participants’ 1-trial place learning performance on the virtual DMP task is strongly associated with theta oscillations in the medial temporal lobe (including the hippocampus) (Bauer and Bast 2020). Overall, the findings reviewed above suggest that the DMP paradigm is a more sensitive assay of hippocampus-dependent navigation than incremental place learning paradigms, as good performance on the DMP task may absolutely require the hippocampus, with extra-hippocampal mechanisms unable to sustain such learning (Bast 2007). In the following sections, we will review some RL approaches that link spatial representations in the hippocampus with successful navigation. We will focus on the performance and limitations of these methods in accounting for navigation based on rapid, one-trial, hippocampal place learning, especially as assessed by the watermaze and virtual DMP tasks in rodents and humans, respectively.

## 3 Reinforcement Learning for spatial navigation

Typical RL problems involve four components: states, actions, values and policy (Sutton and Barto 2018), as shown in Fig.2a. In spatial navigation, states usually represent the agent location, but can be extended to describe more abstract concepts such as contexts or stimuli (Sutton and Barto 2018). Actions are decisions to transition between states (*i.e*. a decision to move from one location to another). Values quantify the mean expected reward to be obtained under a given state or action. Rewards are scalars usually given at certain spatial locations, mimicking goal locations in navigation tasks. The value function can be a function of (*i.e*. dependent upon) the state alone, in which case it refers to the discounted total amount of reward that an agent can expect to receive in the future from a current state *s* at time t. Alternatively, the value can also refer to the state-action pair, in which case it refers to the value of taking a particular action at a certain state. The value function is given by

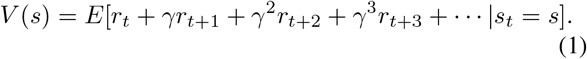

**Figure 2.**
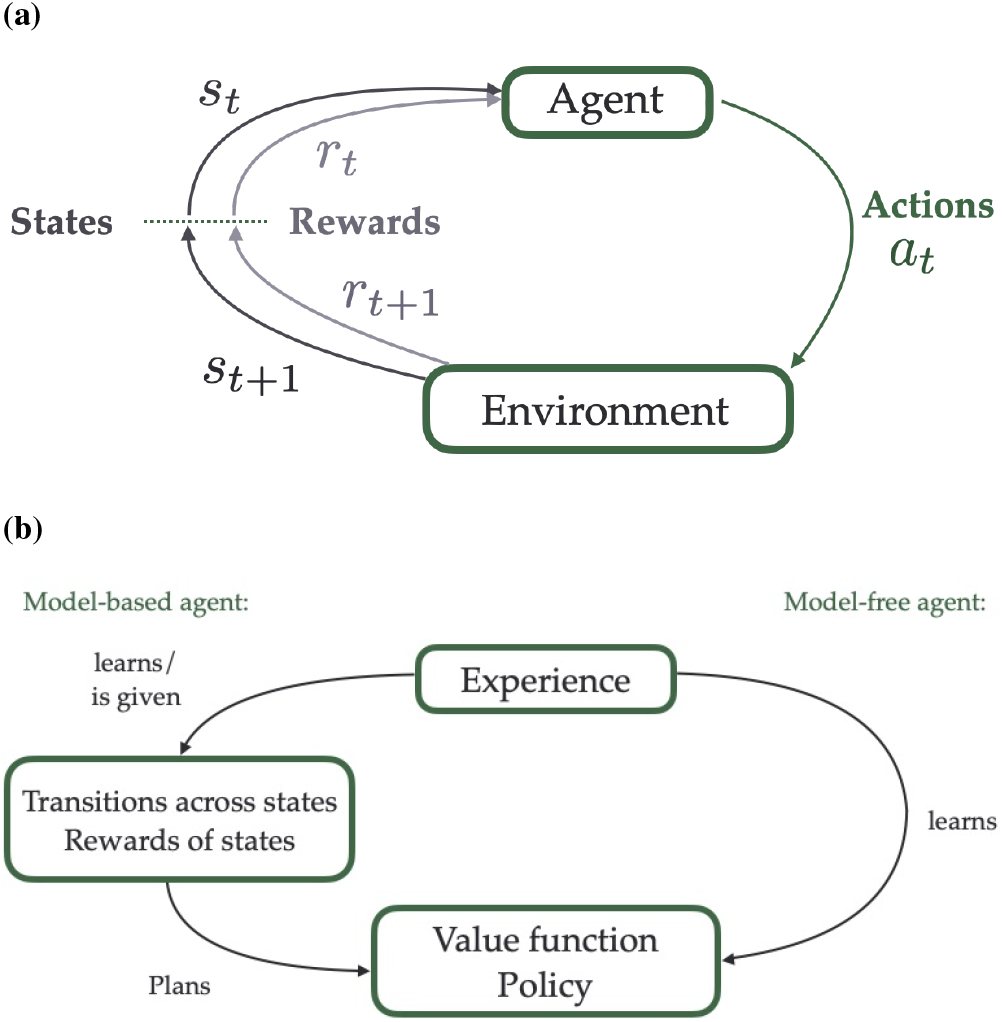
Basic principles of Reinforcement Learning (RL). (a) Key components of RL models. An agent in the state *s_t_* (which in spatial context often corresponds to a specific location in the environment) associated with the reward *r_t_* takes the action *a_t_* to move from one state to another within its environment. Depending on the available routes and on the rewards in the environment, this action leads to the reception of a potential reward *r*_*t*+1_ in the subsequent state (or location) *s*_*t*+1_. (b) Model-free versus model-based approaches in RL. In model-free RL (right), an agent learns the values of the states on the fly, *i.e*. by trial and error, and adjusts its behaviour accordingly (in order to maximise its expected rewards). In model-based RL (left), the agent learns or is given the transition probabilities between states within an environment and the rewards associated with the states (a “model” of the environment), and uses this information to plan ahead and select the most successful trajectory.

In equation (1), the value of state *s, V*(*s*), is computed by summing all future rewards *r_j_* that will be received at time *j, j* ⩾ 0 discounted by a factor *γ*, which quantifies the extent to which immediate rewards are favoured compared to delayed ones of the same magnitude. *E*[·] refers to the expectation, which sums the possible rewards depending on their associated probability. A policy is a probability distribution over the set of possible actions. This defines which actions are more likely to be chosen in a certain location, and has to be learned in order to maximise the *value function* (*i.e*. to maximise the expected amount of reward).

Mathematically, the problem can be represented as a Markov decision process, equipped with transition probabilities between states that shapes the way actions change states, and a reward function that maps states to reward (see, for example,Howard (1960) for more details). In a spatial navigation context, the transition probabilities typically depend on the spatial structure of the environment, and the latter can also vary temporally in certain contexts, as routes might open or close on particular occasions, for example. The probabilities of transition between states/locations in an environment and the rewards available at every location form a *model* of the environment.

In model-free reinforcement learning (Fig.2b, right), the *model* is unknown and the agent must discover its environment, and associated rewards, and learn how to optimise behaviour on the fly, through trial and error. Conversely, in model-based RL approaches (Fig.2b, left), the agent has access to the *model*, from which a tree of possible chains of actions and states can be built and used for planning. In this way the best possible chain of actions can be defined, for example using Dynamic Programming, which selects at every location the optimal action, using one-step transition probabilities (Sutton and Barto 2018).

We can assess whether humans and animals use model-free or model-based strategies by comparing their performance to both types of agent on a two-step decision task (Da Silva and Hare 2019; Miller et al. 2017; Daw et al. 2011). In this task, participants first choose between 2 states, both of which afterwards lead to final states with different probabilities, making one transition “rare” and the other “common”. The final states have unbalanced reward probability distributions (see *e.g* Fig. 1.a. in Miller et al. (2017) for a diagram of the task). Investigating how subjects adjust to rare transition outcomes indicates whether they have access to the *model* or not. A model-free agent will adjust its behaviour only based on the outcome, whereas a model-based agent will adjust also according to the probability of this transition. Both humans and animals show behavioural correlates of model-based and model-free agents (Yin and Knowlton 2006;Keramati et al. 2011;Gershman 2017;Daw et al. 2011;Miller et al. 2017).

Model-based approaches require the calculation and storage of the transition probability matrix and tree-search computations (Huys et al. 2013). As the number of states can be very high, depending on the complexity of the problem and the precision required, model-based methods are usually computationally costly (Huys et al. 2013). However, as they contain exhaustive information about the available routes between states, they are more flexible towards changing goal locations than model-free approaches (Keramati et al.2011). A study of spatial navigation in human participants showed that, although paths to the goal were shorter, choice times were higher in trials when the behaviour matches that of a model-based agent compared to trials where it matches that of a model-free agent (Anggraini et al. 2018). In studies involving rats in a T-maze, Vicarious Trial and Error (VTE) behaviour, short pauses that rats make at decision points, tend to get shorter with repetitive exposure to the same goal location (Redish 2016). Experimental studies suggest that VTE behaviour reflects simulations of scenarios of future trajectories in order to make a decision (Redish 2016), which would correspond to a model-based approach of task solving (Pezzulo et al. 2017;Penny et al. 2013). This suggests that model-based strategies require more processing time than model-free strategies, which is thought to represent “planning” time (Keramati et al. 2011).

The control of behaviour could be coordinated between model-free and model-based systems, either depending on uncertainty (Daw et al. 2005), depending on a trade-off between the cost of engaging in complex computations and the associated improvement in the value of decisions (Pezzulo et al. 2013), or depending on how well the different systems perform on a task (Dollé et al. 2018). Moreover, the state and action representations that enable solution of a task seem to dynamically evolve on the timescale of learning in order to adjust to the task requirements (Dezfouli and Balleine 2019). When the task gives an illusion of determinism - for example when the task is overtrained - the neural representations and the behaviour shift from model-based, purposeful, behaviour, to habitual behaviour, which is faster but less flexible to any change (Smith and Graybiel 2013). When the situation is inherently stochastic, for example when the task evolves to incorporate more steps and complexity, the neural representations and behaviour evolve to incorporate the multi-step dependencies, and simultaneously prune the tree of possible outcomes depending on the most likely scenarios (Tomov et al. 2020). The adaption of neural representations to task demand suggests that a continuum of state and action representations for behavioural control, between the two extremes of modelbased and model-free systems, exists in the brain (Dezfouli and Balleine 2019).

The rapidity with which rats adjust to a changing goal location in the DMP task (one exposure only, as discussed in section 2) indicates that some kind of “model” is being used to enable adaptive route selection. In fact, a model-based approach has recently been proposed to solve a range of spatial learning tasks including the watermaze DMP task (Dollé et al. 2018). Dollé et al. (2018) investigated the interaction between model-free and model-based strategies using a model-free controller to gate interactions between the two. The controller learns to select one of the two strategies depending on the reward in the current task. The model reproduced the flexibility towards new goal locations in the watermaze DMP task, through the gating mechanism, which switched to the model-based strategy for this particular task.

Generally, current model-based approaches, including the one proposed by Dollé et al. (2018), have several limitations in accounting for watermaze DMP task performance in a neuropsychologically realistics way. First, unlike what would be expected based on model-based mechanisms, rats do not reach optimal trajectories in the DMP task, as reflected by the observation that escape latencies on trials 2 to 4 remain higher than would typically be observed following incremental place learning in the watermaze (compare Morris et al. (1986), Steele and Morris (1999) and Bast et al. (2009)) and than would be expected from a model-based agent (Sutton and Barto 2018). Findings in humans by Anggraini et al. (2018) suggest that participants that used more model-based approaches more often took the shortest path to goal locations. Second, classical modelbased approaches are currently mostly implemented in discrete state space, although they can be approximated to continuous spaces (Jong and Stone 2007). The size of the graph to model the environment, requiring a high number of states for fine discretisation to mimic continuity, and the width and depth of the trees to search at possible decision points (for example, at the start location) that would be required to account for such behaviours (every possible trajectories) suggest that a long planning time would be required at a start of every trial (Keramati et al. 2011), which does not fit with the behaviour of rats and human participants, who take off virtually immediately at the start of the second trial to the new goal location on the DMP task in the watermaze and virtual maze, respectively (Steele and Morris 1999; Buckley and Bast 2018). Dollé et al. (2018) overcome this problem using an approximation: the chosen trajectory between the current position and the goal location is in fact the trajectory between their respective closest nodes within the graph. Overall, this suggests that the control in the watermaze DMP task cannot be explained by a model-free RL mechanism alone, but also is unlikely to be fully model-based.

Lying in between model-free and model-based approaches, the Successor Representation (SR) (Dayan 1993) enables more flexibility than model-free computational approaches, but without the heavy computational requirement of a model-based agent (Ducarouge and Sigaud 2017). In the SR, the link between two states depends on how many times the agent can expect to visit one state when starting from the other in the future. It is therefore a predictive representation of space occupancy. Properties of place cell firing, such as shaping of the activity profile by changes in the environment, have been linked to key features of the SR (Stachenfeld et al.2017;Gershman 2018). The SR can be computed from the transition probability matrix, but also can be computed by online learning, usually through counting the occupancy of states (Dayan 1993;Russek et al. 2017). Connecting place cells, using this representation, within an attractor network allows one to generate trajectories from any starting location to any goal location within a maze (Corneil and Gerstner 2015). The SR can be adapted to a continuous state representation (Jong and Stone 2007; Barreto et al. 2017). However, the SR still represents a complex state representation, since the size of the SR matrix is similar to the size of the model-based representation. In the following section, we present two minimal extensions to a model-free architecture that enable flexibility. We will provide an in depth discussion of the underlying computations and of their possible neurobiological substrates.

## 4 A model-free agent using an actor-critic architecture

### 4.1 An actor critic network model for incremental learning

#### 4.1.1 Learning through temporal difference error

Temporal difference (TD) learning refers to a class of model-free RL methods, that improve the estimate of the value function using successive comparison of its value, called the TD error (this approach is commonly referred to as “bootstrapping”). In traditional TD learning, the agent follows a fixed policy and discovers how good this policy is through the computation of the value function (Sutton and Barto 2018). Conversely, in an actor-critic learning architecture, an agent explores the environment and progressively forms a policy that maximises expected reward using a TD error. Interactions with the environment allow simultaneous sampling of the rewards (obtained at certain locations) and of the effect of the policy (when there is no reward, the effect of a policy can be judged depending on the difference in value between two consecutive locations), thereby appraising predictions of values and actions, so that both can be updated accordingly. The “actor” refers to the part of the architecture that learns and executes the policy, and the “critic” to the part that estimates the value function (Fig.3a).

**Figure 3.**
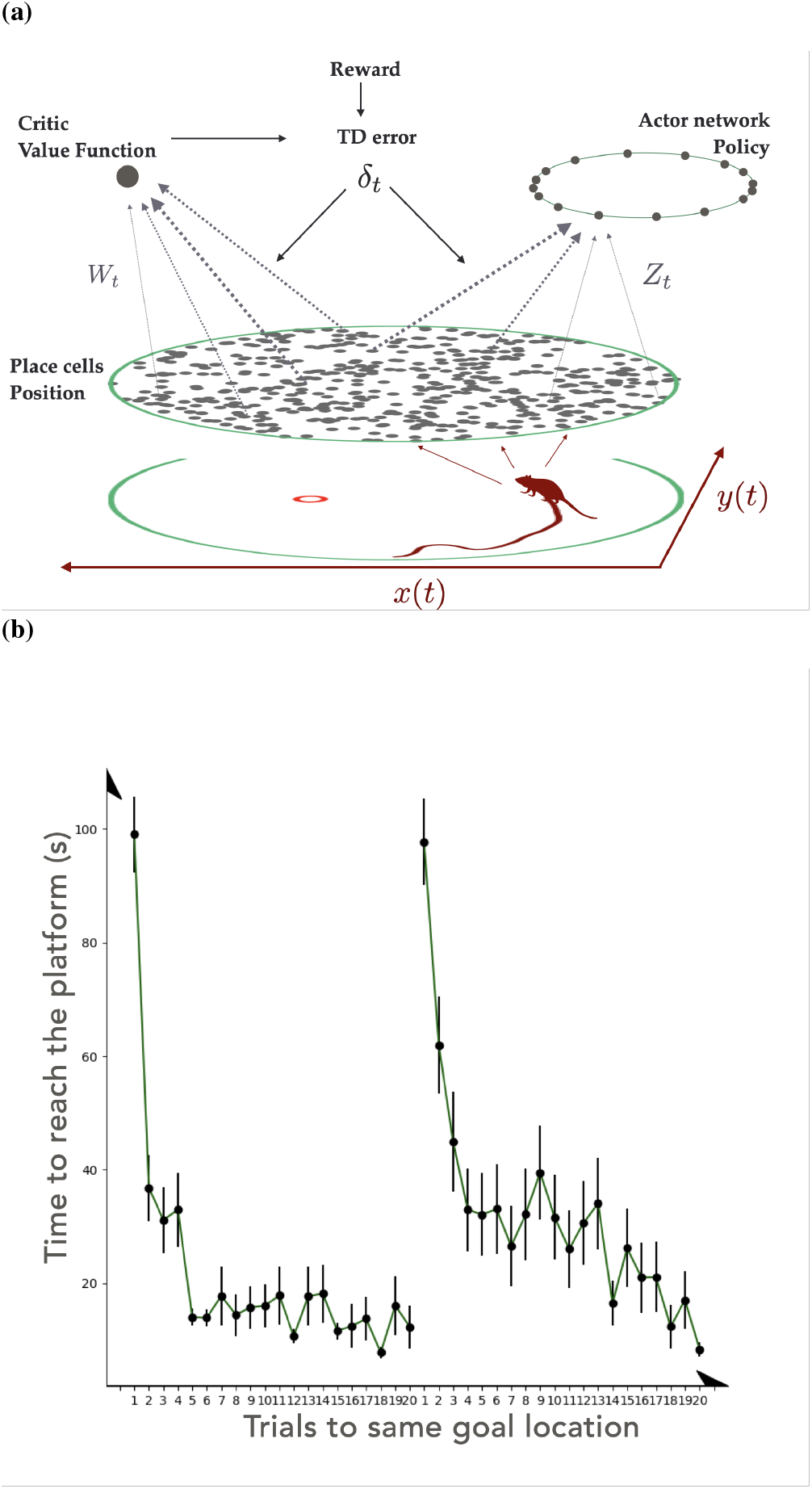
a) Classical actor-critic architecture for a temporaldifference (TD) agent learning to solve a spatial navigation problem in the watermaze, as proposed by Foster et al. (2000). The state (location of the agent (*x*(*t*), *y*(*t*))) is encoded within a neural network (in this case, the units mimic place cells in the hippocampus). State information is fed into an action network, which computes the best direction to go next, and to a critic network that computes the value of the states encountered. The difference in critic activity along with the reception or not of the reward (given at the goal location) are used to compute the TD error *δ_t_*, such that successful moves (that lead to a positive TD error) are more likely to be taken again in the future, and less likely otherwise. Simultaneously, the critic’s estimation of the value function is adjusted in order to be more accurate. These updates occur through changing the critic and actor weights, respectively *W_t_* and *Z_t_*. The goal location, marked as a circle within the maze, is the only location in which a reward is given. b) Performance of the agent, obtained by implementing the model from Foster et al. (2000). The time that the agent requires to get to the goal (“Latencies”, vertical axis) reduces with trials (horizontal axis) and reaches almost a minimum (after trial 5). When the goal changes (on trial 20), the agent takes a very long time to adapt to this new goal location.

Foster et al. (2000) proposed an actor-critic framework to perform spatial learning in a watermaze equivalent (Fig.3a). The agent’s location is represented through a network of units mimicking hippocampal place cells, which have a Gaussian type of activity around their preferred location (the further the agent is from the place cell’s preferred location, the less active the unit will be). Each place cell projects to an actor network and a critic unit, or network in other related models, *e.g*. in Frémaux et al. (2013), through plastic connections (respectively denoted *Z_t_* and *W_t_* in Fig.3a).

The actor network activity defines which action is optimal, with each unit in the network coding for motion in a particular direction, together covering a 360°angle. In Foster et al. (2000) this angular direction is quantised, whereas in Frémaux et al. (2013), using a more detailed spiking neuron model, the action network codes for continuous movement directions (although finely quantised in numerical simulations). The action is chosen according to the activity of the actor, so that directions corresponding to more active cells are prioritised. In Foster et al. (2000), the action corresponding to the maximum of a softmax probability distribution of the activities is selected. In Frémaux et al. (2013), the activities of the units of the network are used as weights to sum all possible directions, giving rise to a mean vector that defines the chosen direction. If the motion leads to the goal location, a reward is obtained. In the case of computational models of the watermaze task, the reward is only delivered when the agent is within a small circle representing the platform (see Fig.3a). This reward information, encoded within the environment is used along with the difference in the successive critic activities to compute the TD error *δ_t_*, via

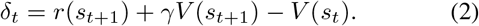

In equation 2, the reward observed after a move *r*(*s*_*t*+1_) along with the discounted value of the new state *γV*(*s*_*t*+1_) is compared to the prior prediction *V*(*s_t_*). The TD error contains two pieces of information: it reflects both how good is the decision that was just taken, and the accuracy of the critic at estimating the value of the state.

To reduce the TD error, the model simultaneously updates the connections weights *Z_t_* and *W_t_* via defined learning rules in order to improve the actor and the critic. The learning rule updates the connection weights according to the current error and the place cell activity, such that the probability of taking the same decision in the current state/location increases if it leads to a new position that has a higher value, and decreases otherwise (see Doya (2000) for example learning rules). Using the model proposed by Foster et al. (2000), we can reproduce their finding that within a few trials, the agent reaches the goal using an almost optimal path, reflected by low latencies to reach the goal, in a watermaze-like environment (Fig.3b). The full model can be found in appendix A.

#### 4.1.2 Important variables for learning: discount factor for value propagation, and spatial scale of place cells representations for experience generalisation

The previous section illustrated how place cells can be integrated into a network for spatial navigation through the association of values and actions within a RL framework. A benefit of this approach is that i) it allows for relatively fast learning, within only a few trials agents reach “short” average escape latencies and ii) the agent obtains information about which action could lead to the goal from variable and distal start positions. These two properties rely on two major components that enable learning and influence the learning speed (*i.e*., how fast latencies reduce).

First, the TD error allows to “back-propagate” value information from successive states, with the speed of this backpropagation modulated by the discount factor *γ*. The update of the state value *V*(*s_t_*) and of the policy at state *s_t_*, after having moved to the new state *s*_*t*+1_, depends on the TD error. Let us consider the latter, defined by equation (2): the TD error is the difference between the received reward and the discounted value at the next state (given by *r*(*s*_*t*+1_) + *γV*(*s*_*t*+1_), the first two terms in Eq.(2)), and the value at the current state (*V*(*s_t_*), the last term in Eq.(2)). Note that the update of the state value is performed “in the future” for “the past”: the value underlying the decision taken at the time *t* will only be updated at the time *t* + 1. Moreover, the extent to which the future is taken into account is modulated by the parameter *γ*.

Let us consider the extreme cases. If *γ* = 0, the only update takes place when the reward is found, and the location updated is the one immediately prior to the reward. All the other locations, which do not precede the reward reception, will never be associated with a non-zero value. If *γ* = 1, the state value *V*(*s_t_*) will be updated until it is equal to *V*(*s*_*t*+1_). This leads to a constant value function over the maze (*i.e*. all locations have the same value). In both cases, the value function being uniform, the actor computes all actions as equally good (except very near the goal in the case where *γ* = 0), as it moves through the environment. Therefore, only intermediate values of *γ* allow the model to learn, and its value defines how fast it learns. Ideally, one wants to adjust the discount factor *γ* in order to maximise the slope of the value function, so that the policy is “concentrated” on the optimal choice, and to obtain a uniform slope across the space, as this guarantees that the agent has good information on which to base its decision from any location within the environment.

Second, the spatial scale of the place cell representation, determined by the width of these neurons’ place fields, strongly affects speed and precision of place learning. The state representation through place cells enables the generalisation of learning from a single experience across states, *i.e*. to update information on many locations based on the experience within one particular location. Every update amends the value and policy for all states depending on the current place cell activity, such that more distal locations are less concerned by the update than proximal ones. The spatial reach of a particular update increases with the width of the place cell activity profiles. This process speeds up learning, because when the agent encounters a location with similar place cell representation to those already encountered, the prior experiences have already shaped the current policy and value function and can be used to inform subsequent actions.

Let us consider the extreme cases. If the width is very small, the agent cannot generalise enough from experience, and this considerably slows down learning, as the agent must comprehensively search the environment in order to learn. At the opposite extreme, if the activity profile is very wide, generalisation occurs where it is not appropriate: for example at opposite ends of the goal, when the best actions to choose would be opposite to each other, as at the North end it would be best to go South, whereas at the South end it would be best to go North. The optimal width, therefore, lies in a trade-off between speed of learning and precision of knowledge: it should be scaled to the size of the environment in order to speed up learning, and is constrained by the size of the goal. Optimising these parameters allows one to reduce the number of trials to obtain good performance. Appendix B shows how changing these parameters affects learning.

Along the hippocampal longitudinal axis, places are represented over a continuous range of spatial scales, with the width of place cell activity profiles gradually increasing from the dorsal (also known as septal) towards the ventral (also known as temporal) end of the hippocampus in rats (Kjelstrup et al. 2008). A recent RL model suggests that smaller scales of representation would support the generation of optimal path length, whereas larger scales would enable faster learning, defining a trade-off between path optimality and speed of learning (Scleidorovich et al. 2020). In Fig. 6 of appendix B, for the actor-critic model, we also see that a wide activity profile of place cells leads to suboptimal routes, characterised by escape latencies that stay high.

Bast et al. (2009) found that the intermediate hippocampus is critical to maintain DMP performance in the watermaze, particularly search preference. Moreover, the trajectories used by rats in the watermaze DMP task are suboptimal, *i.e*. path lengths are higher, compared to the incremental learning task (compare results in Morris et al. (1990), Steele and Morris (1999) and Bast et al. (2009)). These findings may partly reflect that place neurons in the intermediate hippocampus, which show place fields of an intermediate width and, thereby, may deliver a trade off between fast and precise learning, are particularly important for navigation performance during the first few trials of learning a new place, as on the DMP task. Another potential explanation for the importance of the intermediate hippocampus is that this region combines sufficiently accurate place representations, provided by place cells with intermediate-width place fields, with strong connectivity to prefrontal and subcortical sites that support use of these place representations for navigation (Bast et al. 2009; Bast 2011), including striatal RL mechanisms (Humphries and Prescott 2010). With incremental learning of a goal location, spatial navigation can become more precise, with path lengths getting shorter and search preference values increasing (Bast et al. 2009). Interestingly, the model by Scleidorovich et al. (2020) suggests that such precise incremental place learning performance may be particularly dependent on narrow place fields, which are shown by place cells in the dorsal hippocampus (Kjelstrup et al. 2008). This may help to understand why incremental place learning performance has been found to be particularly dependent on the dorsal hippocampus (Moser et al. 1995).

#### 4.1.3 Using eligibility traces allows to update past decisions according to the current experience

The particular actor-critic implementation proposed by Foster et al. (2000) and described above involves a one-step update: only the value and policy of the state that the agent just left are updated. The weight associated to the value of the following state for any update depends on γ, as described in the previous section. However, past decisions sometimes affect current situations, and one-step updates only improve the last choice and estimate. This can be addressed by incorporating past decisions when performing the current update, weighted according to an *eligibility trace* (Sutton and Barto 2018). Eligibility traces keep a record of how much past decisions influence the current situation. This makes possible to update value functions and policies from previous states of the same trajectory according to the current outcome.

The most commonly known example of the use of eligibility traces is TD(λ) learning, which updates the value and policy of previous states within the same trajectory according to the outcome observed after a certain subsequent period, weighted by a decay rate λ. λ refers to how far in the past the current situation affects previous states’ value and policy. One extreme is TD(0) in which the only step updated is the state the agent just left, as described in section 4.1. When λ increases toward 1, more events within the trajectory are taken into account, and the method becomes more reminiscent of a Monte-Carlo approach (Sutton and Barto 2018), where all the states and actions encountered during the trial are updated at every step. In (Scleidorovich et al. 2020), the authors show that the optimal value of λ depends on the width of the place cell activity distributions: for wide place fields, adding eligibility traces does not speed up learning much (*i.e*. reducing the number of trials needed to reach asymptotic performance), but it does for narrower place fields.

#### 4.1.4 Striatal and dopaminergic mechanisms as candidate substrates for the actor and critic components

In the RL literature, the ventral striatum is often considered to play the role of the “critic” (Humphries and Prescott 2010; Khamassi and Humphries 2012; O’Doherty et al. 2004; van Der Meer and Redish 2011). The firing of neurons in the ventral striatum ramps up when rats approach a goal location in a T-maze (van Der Meer and Redish 2011), consistent with the critic activity representing the value function in actor-critic models of spatial navigation (Foster et al. 2000; Frémaux et al. 2013). Striatal activity also correlates with action selection (Kimchi and Laubach 2009), and with action specific reward values (Roesch et al. 2009;Samejima et al. 2005).

In line with the architecture proposed in the model by Foster et al. (2000) (Fig.3a), there are hippocampal projections to the ventral and medial dorsal striatum (Groenewegen et al. 1987; Humphries and Prescott 2010). Studies combining watermaze testing with manipulations of ventral and medial dorsal striatum support the notion that these regions are required for spatial navigation. Lesions of the ventral striatum (Annett et al. 1989) and of the medial dorsal striatum (Devan and White 1999) have been reported to impair spatial navigation on the incremental place learning task. In addition, crossed unilateral lesions disconnecting hippocampus and medial dorsal striatum also impair incremental place learning performance, suggesting hippocampo-striatal interactions are required (Devan and White 1999). To our knowledge, it has not been tested experimentally if there is a dichotomy between “actor” and “critic”. The experimental evidence outlined above is consistent with both actor and critic roles of the striatum (van Der Meer and Redish 2011), but whether distinct or the same striatal neurons or regions act as actor and critic needs to be addressed.

Li and Daw (2011) address a related dichotomy in a study on human participants who have to choose between two arms associated with reward probabilities on a bandit task. The participants are given the outcomes of their decision after their choice: namely how much they win and how much they would have won if they had selected the other arm. Li and Daw (2011) compare two ways of updating the weights which determine which arm to choose: one compares the reward to the predicted value (“value update”), and the other one compares the reward to the forgone reward (“policy update”). They show that striatal BOLD activity correlates more with “policy” than “value” update, and correlates positively with the chosen reward, and negatively with the reward that was not chosen. They also show correlation with a value-based decision variable, the difference between the action value of the chosen and the arm not chosen. The translation of their analysis to spatial navigation in the watermaze is not straightforward. First, in the two armed bandit task, there are no states, but only actions. Although the design of the analysis by Li and Daw (2011) allows one to disentangle the rewards from the predicted values, it does not allow one to separate action from state value in a spatial navigation context, if it makes sense at all to separate the two. In spatial navigation, as states can be passed through to reach any goal, it seems to be more efficient not to separate actions and values. However, it is interesting to see an experimental setup aimed at testing such a dichotomoty. Perhaps, an architecture such as SARSA (State–Action–Reward–State–Action, Sutton and Barto (2018)), in which the values of a state action pair are learned instead of the values of states only, could be considered, as it unites the actor and critic computation within the same network.

Phasic dopamine release from dopaminergic midbrain projections to the striatum has long been suggested to reflect reward prediction errors (Schultz et al. 1997; Glimcher 2011), which correspond to the TD error in the model in Fig.3a, and dopamine release in the striatum shapes action selection (Gerfen and Surmeier 2011;Morris et al. 2010;Humphries et al. 2012). Direct optogenetic manipulation of striatal neurons expressing dopamine receptors modified decisions (Tai et al. 2012), consistent with the actor activity in actor-critic models of spatial navigation (Foster et al. 2000; Frémaux et al. 2013). Moreover, 6-hydroxydopamine lesions to the striatum, depleting striatal dopamine (and, although to a lesser extent, also dopamine in other regions, including hippocampus) impaired spatial navigation on the incremental place learning task in the watermaze (Braun et al. 2012). These findings suggest that aspects of the dopaminergic influence on striatal activity could be consistent with the modulation of action selection by the TD errors in an actor-critic architecture.

However, although there is long term potentiation (LTP)-like synaptic plasticity at hippocampo-ventral striatal connections, consistent with the plastic connections between the place cell network and the critic and actor in the model by Foster et al. (2000), a recent study failed to provide evidence that this plasticity depends on dopamine (LeGates et al. 2018). Absence of dopamine modulation of hippocampo-striatal plasticity contrasts with the suggested modulation of connections between place cell representations and the critic and actor components by the TD error signal in the RL model. Thus, currently available evidence fails to support one key feature of the architecture described in section 4.1 (Foster et al. 2000).

#### 4.1.5 Requirement of hippocampal plasticity

In the implementation of the model described above, plasticity takes place within the feedforward connections from the place cell network modelling the hippocampus and the actor and critic networks that, as discussed above, could correspond to parts of the striatum. The model does not capture the finding that hippocampal NMDA receptor-dependent plasticity is required for incremental place learning in the watermaze if rats have not been pretrained on the task (Morris et al. 1986, 1989; Nakazawa et al. 2004).

#### 4.1.6 The agent is less flexible than animals in adapting to changing goal locations

When the goal changes (on trial 20 in Fig.3b), the agent takes many trials to adapt and takes more than 10 trials to reach asymptotic performance levels (also see Fig.4a in Foster et al. (2000)). The high accuracy, but limited flexibility with overtraining, are well known features of TD RL methods (e.g. discussed in Sutton and Barto (2018); Gershman (2017); Gershman et al. (2014); Botvinick et al. (2019)). These cached methods have been proposed to account for the progressive development of habitual behaviours (Balleine 2019). TD learning is essentially an implementation of Thorndike’s law of effect (Thorndike 1927), which increases the probability of reproducing an action if it is positively rewarded. In the RL model discussed above (Fig.3a), a particular location, represented by activities of place cells with overlapping place fields, is associated with only one “preferred action”, due to the unique weights that need to be fully relearned when the goal changes. Therefore, the way actions and states are linked only allows navigation to one specific goal location.

**Figure 4.**
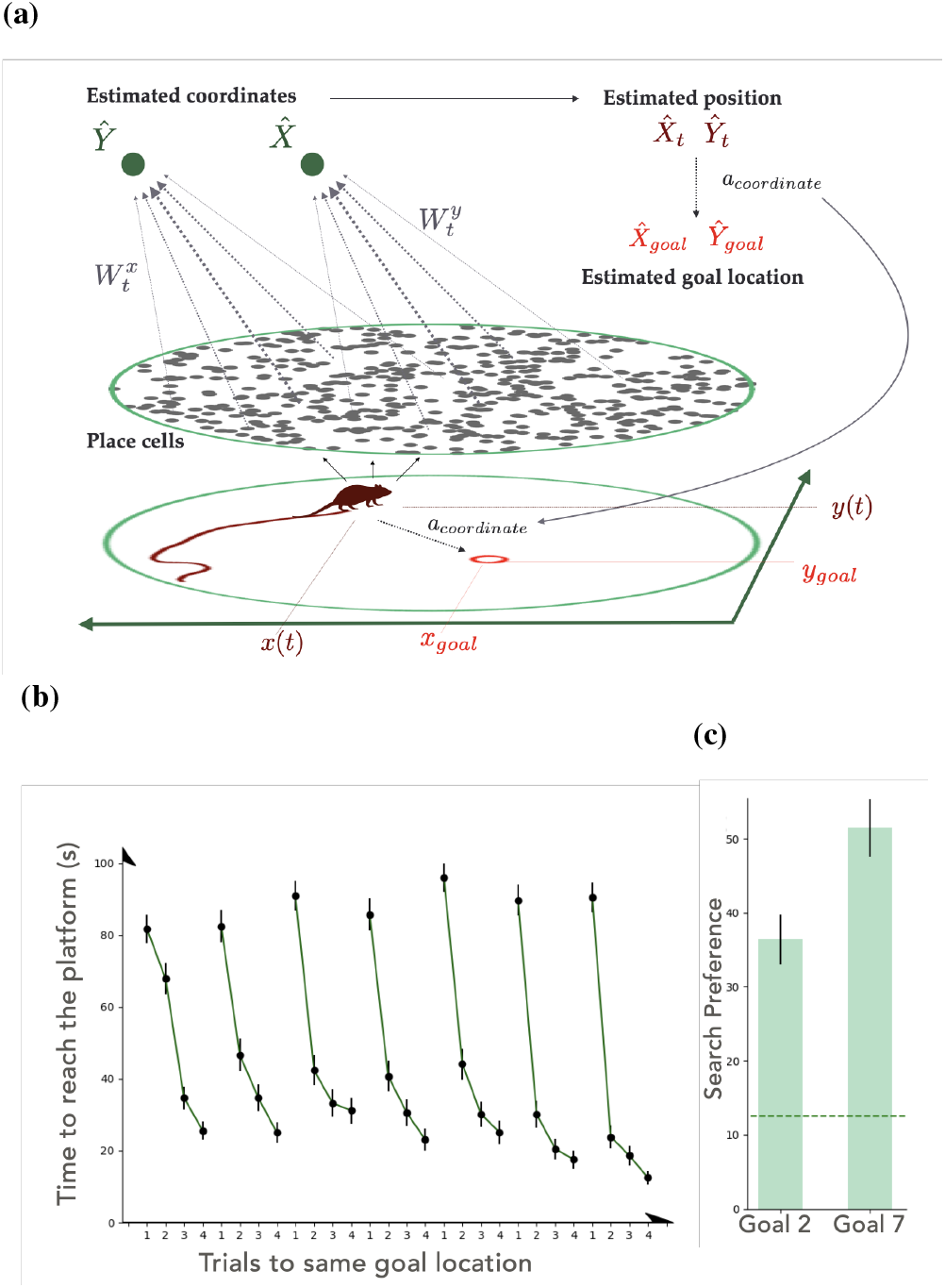
(a) Architecture of the coordinate-based navigation system, which was added to the actor-critic system shown in Fig. 3a to reproduce accurate spatial navigation based on one-trial place learning, as observed in the watermaze DMP task (Foster et al. 2000). Place cells are linked to coordinate estimators through plastic connections 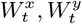. The estimated coordinates 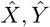 are used to compare the estimated location of the goal 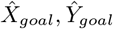 to the agent estimated location 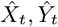 in order to form a vector towards the goal, that is being followed when choosing the “coordinate action” *a_coord_*. The new action *a_coord_* is integrated into the actor network described in Fig.3a. (b,c) Performance of the extended model using coordinate-based navigation. (b) Escape latencies of the agent when the goal location is changed every four trials, mimicking the watermaze DMP task. (c) “Search preference” for the area surrounding the goal location, as reflected by the percentage of time the agent spends in an area centered around the goal location when the second trial to a new goal location is run as probe trial, with the goal removed (stippled line indicates percentage of time spent in the correct zone by chance, *i.e*. if the agent had no preference for any particular area), computed for the second and the seventh goal locations. One-trial learning of the new goal location is reflected by the marked latency reduction from trial 1 to trial 2 to a new goal location (without interference between successive goal locations), and by the marked search preference for the new goal location when trial 2 is run as probe. The data in (b) were obtained by computing the model in (Foster et al. 2000), and the data in (c) by adapting the model in order to reproduce search preference measures when trial 2 was run as a probe trial. The increase in search preference observed between the second and seventh goal location is addressed in section 4.2.1.

The model produces a general control mechanism, that, in this example, makes it possible to generate trajectories to a particular goal location. This control mechanism could be integrated within an architecture that allows more flexibility, for example the goal location may be represented by different means than a unique value function computed via slow and incremental steps from visits to single goal locations. The next section considers how the RL model of Fig.3a can be used, along with a uniform representation of both the agent and the goal location, to reproduce the flexibility shown by rats and humans towards changing goal locations in the watermaze DMP task.

### 4.2 Map-like representation of locations for goal-directed trajectories

The RL architecture discussed above (Fig.3a) cannot reproduce rapid learning of a new location as observed in the DMP watermaze task, but instead there is substantial interference between successive goal locations, with latencies increasing across goal locations, and only a gradual small decrease in latencies across the four trials to the same goal location (see Foster et al. (2000), Fig. 4b). To reproduce flexible spatial navigation based on one-trial place learning as observed on the DMP task, Foster et al. (2000) proposed to incorporate a coordinate system into their original actor critic architecture (Fig.4a). This coordinate system is composed of two additional cells 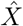 and 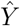 that learn to estimate the real coordinates *x* and *y* throughout the maze. These cells receive input from the place cell network through plastic connections 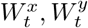. The connections evolve dependent on a TD error that represents the difference between the displacement estimated from the coordinate cells and the real displacement of the agent. The weights between place cells and the coordinate cells are reduced if the estimated displacement is higher than the actual displacement, and increased if it is lower, so that the estimated coordinates progressively become consistent with the real coordinates (see Fig.8 in Appendix D).

Foster et al. (2000) added an additional action to the set of actions already available. Instead of defining movement in specific allocentric directions, as the other action cells do, going North, East, etc…, that we will refer to as “allocentric direction cells”, the coordinate action cell *a_coord_* points the agent in the direction of the estimated goal location. The estimates of *x* and *y* are used to compare the agent’s estimated position 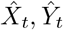 to the estimated goal location 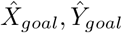 (which is stored after the first trial of every day) in order to form a vector leading to the estimated goal location (Fig.4a).

The agent very quickly adapts to new goal locations, reproducing performance similar to rats and humans on the watermaze (Fig. 1) and virtual (Buckley and Bast 2018) DMP task, respectively. Using the model by Foster et al. (2000), we can replicate their finding that the model reproduces the characteristic pattern of latencies shown by rats and human on the DMP task, *i.e*. a marked reduction from trial 1 to 2 to a new goal location and no interference between successive goal location (Fig. 4b). Moreover, extending the findings of Foster et al. (2000), we find that the model also reproduces markedly above-chance search preference for the vicinity of the goal location when trial 2 to a new goal location is run as probe trial where the platform is removed (Fig. 4c).

#### 4.2.1 Limitations of the model in reproducing DMP behaviour in rats and humans

The “coordinate” approach relies on computational “tricks” that are are required to make the approach work, but for which plausible neurobiological substrates remain to be identified. Early in training, movement of the agent is based on activity of the “allocentric direction cells”, which are used to lead the exploration of the environment. This exploratory phase allows learning of the coordinates. As the estimated coordinates 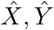 become more consistent with the real coordinates, the coordinate action *a_coord_* becomes more reliable, as it will always lead the agent in the direction of the goal. During the first trial to a new goal location, the coordinate action cell encodes random displacement, until the goal is found and its estimated location is stored. During this trial, the coordinate action is not reinforced, a trick that prevents its devaluation. On the subsequent trials, the coordinate action encodes the displacement towards the stored estimated goal location (as described before) and is reinforced. Therefore, the probability of choosing the coordinate action gradually becomes one, and it comes to be the only action followed.

One consequence of this is that, unlike in rats and people, the agent’s performance both in terms of latency reduction and in terms of search preference gradually improves across successive new goal locations (see Fig.4b and 4c). The gradual improvement of latency reductions and search preferences contrasts with behaviour shown by rats (Fig.1 and see Fig.3. b. and c. in Bast et al. (2009) for search preference across days) and human participants (Buckley and Bast 2018). More specifically, in the model, latency reductions from trial 1 to 2 gradually increase across successive new goal locations, and the latencies on trial 2 to 4 to a new goal location gradually decrease (Fig.4b). In contrast, rats basically reach asymptotic performance levels with no systematic increases in latency reductions from trial 1 to 2 or decreases in latency values on trial 2 to 4 after a few successive goal locations; in the example shown in Fig.1b, asymptotic performance levels are reached from about day 4. It should also be noted that the overall latency reductions across the first few locations in rats is likely to mainly reflect procedural learning, with rats learning that they cannot escape by climbing the wall of the pool or by diving. Human participants on the virtual DMP task, who do not need to learn the task requirements because they receive task instructions, show virtually asymptotic latency and path lengths values from the first new goal locations, with hardly any improvements across successive goal locations (Buckley and Bast 2018). Moreover, search preference for the correct location substantially increases across successive probe trials in the model (Fig.4c), whereas in rats and humans search preference remains stable across successive new goal locations on the DMP task (Bast et al.2009;Buckley and Bast 2018).

In addition, the random search during trial 1 in the model is inconsistent with the finding that rats on the DMP task (but not human participants, (Buckley and Bast 2018)) tend to go towards the previous goal location on trial 1 to a new goal location (Steele and Morris (1999);Pearce et al. (1998), and our own unpublished observations); in addition, both rats and human participants show systematic search patterns on trial 1 (Buckley and Bast 2018;Gehring et al.2015). The random search in the model during trial 1 leads to consistently and similarly high trial 1 latencies (Fig.4b). In contrast, in rats, trial 1 latencies are more variable (Fig. 1b); this partly reflects procedural learning across the first few new goal locations, which results in reductions of trial 1 latencies, and the different spatial relationship between the start location and the previous and current goal location affecting trial 1 latencies (e.g., if the current goal location lies on the path from the start location to the previous goal location, rats are more likely to “bump” into the current goal location on trial 1, leading to lower trial 1 latencies). The adjustment of the policy when the predicted goal is not encountered is not addressed in the current approach, a point that section 4.3 will address.

#### 4.2.2 The model’s actor-critic component and striatal contributions to DMP performance

Given the association of actor-critic mechanisms with the striatum (Joel et al. 2002; Khamassi and Humphries 2012; O’Doherty et al. 2004; van Der Meer and Redish 2011), the actor-critic component in the model is consistent with our recent findings that the striatum is associated with rapid place learning performance on the DMP task. More specifically, using functional inhibition of the ventral striatum in rats, we have shown that the ventral striatum is required for one-trial place learning performance on the watermaze DMP task (Seaton 2019); moreover, using high-density electroencephalogram (EEG) recordings with source reconstruction in human participants, we found that theta oscillations in a circuit including both temporal lobe and striatum are associated with one-trial place learning performance on the virtual DMP task (Bauer and Bast 2020).

The model suggests that, after a few trials, once the action probability for the coordinate action has reached the value 1, the movement is predefined by following a vector pointing to the goal location. The critic becomes inconsistent, as the action now does not follow the value gradient anymore, and, therefore, there is no control over the behaviour by the TD error. The continued association of the striatum with DMP performance, beyond the first few trials, is consistent with the role of the striatum as the “actor” (van Der Meer and Redish 2011), and the model would suggest that the striatum reads out estimated locations and computes a vector towards the estimated goal location.

#### 4.2.3 Neural substrates of the goal representation and hippocampal plasticity required for rapid learning of new goal locations

Notwithstanding some limitations, findings with this model support the important idea that, embedded within a model-free RL framework, a map-like representation of locations within an environment may allow computations by the agent to produce efficient navigation to new goal locations within as little as one trial. This idea is also present in recently proposed agents that are capable of flexible spatial navigation based on a RL system complemented by path integration mechanisms to derive a grid-like map of the environment (which resembles entorhinal grid cell representations) that can be used to compute trajectories from the agent’s location to the goal (Banino et al. 2018) and has also lead the watermaze DMP task to be solved using a graph search algorithm in Dollé et al. (2018). The findings of goal-vector cells in the bat hippocampus (Sarel et al. 2017) and of “predictive reward place cells” in mouse hippocampus (Gauthier and Tank 2018) support the idea implemented in the model that consistency of representations - unified representations of goals and locations across tasks and environment - could help goal-directed navigation. In particular, egocentric boundary encoding neurons have been found in the striatum of rats, although in the dorso-medial striatum (Hinman et al. 2019). As rats navigate in the watermaze using surrounding cues, these cells could inform striatal navigation in the DMP task (Bicanski and Burgess 2020).

In the extension to the classical TD architecture (Foster et al. 2000), the encounter with a new goal location does not involve a change in place cell representation, and the formation of the memory of the new goal location is not addressed. Experimental evidence suggests that a goal representation could lie within hippocampal representations themselves (Hok et al. 2007;Poucet and Hok 2017; McKenzie et al. 2013; Gauthier and Tank 2018). McKenzie et al. (2013) studied hippocampal CA1 representations during learning of new goal locations in an environment where previous places were already associated with goals. They showed that neurons coding for existing goals would also encode new goal locations, and that these representations progressively separate with repetitive learning of the new goal location, but maintain an overlap of representations between all goal locations. Moreover, Hok et al. (2007) observed an increase in firing rate around goal locations outside of place cells’ main firing field, and Dupret et al. (2010) showed that learning of new goal locations by rats in a food-reinforced dry-land DMP task is associated with an increase in the number of CA1 neurons that have a place field around the goal location. Furthermore, Dupret et al. (2010) showed that both this accumulation of place fields around the goal location and rapid learning of new goal locations is disrupted by systemic NMDA receptor blockade. These findings suggest that goal representation can be embedded within the hippocampus, that new goal locations are represented within similar networks as previous goal locations, and that the hippocampal remapping emerging from new goal locations is linked to behavioural performance and may depend on NMDA-receptor mediated synaptic plasticity.

In line with this suggestion, studies, combining intra-hippocampal infusion of an NMDA receptor antagonist with behavioural testing and electrophysiological measurements of hippocampal LTP, showed that hippocampal NMDA-receptor dependent LTP-like synaptic plasticity is required during trial 1 for rats to learn a new goal location in the watermaze DMP task (Steele and Morris 1999) and in a dry-land DMP task (Bast et al. 2005). LTP-like synaptic plasticity may give rise to changes in place cell representations (Dragoi et al. 2003), which could contribute to changes in hippocampal place cell networks associated with the learning of new goal locations (Dupret et al. 2010).

Map-like representations of locations, integrated within a RL architecture, may be part of neural mechanisms that enable flexibility to changing goal locations in the watermaze DMP task. However, cartesian coordinates are convenient here because the task is implemented within an open field arena, but do not seem to provide a biologically realistic implementation of spatial navigation problems in general. For example, they do not allow navigation in an environment with walls, for which geodesic coordinates would be more appropriate (Gustafson and Daw 2011). Moreover, the approach described here does not address how the goal representation comes about, and the model does not specify how the policy adjusts when the agent does not encounter the predicted goal. The next section describes how a hierarchical architecture can provide a solution to this problem.

### 4.3 Hierarchical control to flexibly adjust to task requirements

The actor-critic approach described in section 4.1 requires many trials to adjust to changes in goal locations partly because there is only one possible association between location and action, which depends upon a particular goal (4.1.6). However, brains are able to perform multiple tasks in the same environment. Those tasks often involve sequential behaviour at multiple timescales (Bouchacourt et al. 2019). Pursuing goals sometimes requires following a sequence of sub-routines, with short-term/interim objectives, themselves divided into elemental skills (Botvinick et al. 2019). Hierarchically organised goal-directed behaviours allow computational RL agents to be more flexible (Dayan and Hinton 1993; Botvinick et al. 2009).

In the watermaze DMP task, rats tend to navigate to the previous goal location on trial 1 with a new goal location (Steele and Morris (1999); Pearce et al. (1998), and our own unpublished observations), and then find out that this remembered goal location is not the current goal. This suggests that preexisting goal networks can flexibly adjust to errors, and are linked to control mechanisms over shorter timescales that allow movement realisation in order to navigate to the new goal location. The critic controls the selection of direction at every time step, finely chosen to mimic the generation of a smooth trajectory. The critic sits at an intermediary level of control: It does not perform the lower level of the control, the motor mechanisms responsible for the generation of the limb movements, but also does not control the choice or retrieval process of the goal that is being pursued. The critic allows progressive decisions in order to reach one particular goal (van Der Meer and Redish 2011).

We hypothesise that the computation of a goal prediction error within a hierarchical architecture could enable flexibility towards changing goal locations. We implemented a hierarchical agent, but the architecture itself does not perform hierarchical learning. In Botvinick (2012) agents are trained to find sub-routines, for example through looking for bottleneck states in a graph. In our implementation, we simply added a layer, which selects which one of the subroutines is most suited for the current situation.

In our implementation of a hierarchical RL architecture to allow for flexible one-trial place learning, we consider familiar goal locations, *i.e*. although the goal location changes every four trials, the goal locations are always chosen from a set of 8 locations where the agent has learned to navigate to the goal during a pretraining period. This contrasts with the most commonly used watermaze DMP procedure where the goal locations are novel (*e.g*., (Steele and Morris 1999; Bast et al. 2009)), although there are also DMP variations where the platform location changes daily, but is always chosen from a limited number of platform locations (Whishaw 1985). Pre-trained, familiar, goal locations are also a feature of delayed-non matching-to-place (DNMP) tasks in the radial maze (e.g., Lee and Kesner (2002),Floresco et al. (1997)), where rats are first pre-trained to learn that food can be found in any of 8 arms (*i.e*., these are familiar goal locations); after this, the rats are required to use a ‘non-matching-to-place’ rule to chose between several open arms during daily test trials, based on whether they found food in the arms during a daily study or sample trial: arms that contained food during the sample trial will not contain food during the test trial and vice versa. Based on work by Schweighofer and Doya (2003), we propose that the agent’s behaviour could be shaped by the chosen policy depending on how confident the agent is about the policy leading to the goal.

The agent learns different policies and value functions using the model described in section 4.1, each of them associated with one of eight possible goal locations presented in the DMP task (which would be at the centre of the eight zones shown in Fig.1c). After multiple trials necessary to learn the actor and critic weights (as presented in section 4.1) for each goal location, the policies and value functions associated with each one of them are stored. We refer to the value and associated policy as a “strategy”. The choice of a strategy depends on a goal prediction error 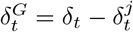 (see Fig.5a). The goal prediction error is used to compute a level of confidence that the agent has in the strategy it follows. When the strategy followed does not lead to the goal, the confidence level decreases, leading to more exploration of the environment until the goal is discovered. Once the goal is discovered, the strategy that minimises the prediction error is selected. Fig.5b represents the latencies of the agent. The agent can quickly adapt to changing goal locations, as reflected by the steep reduction in latencies between trial 1 and 2 of the new goal location.

**Figure 5.**
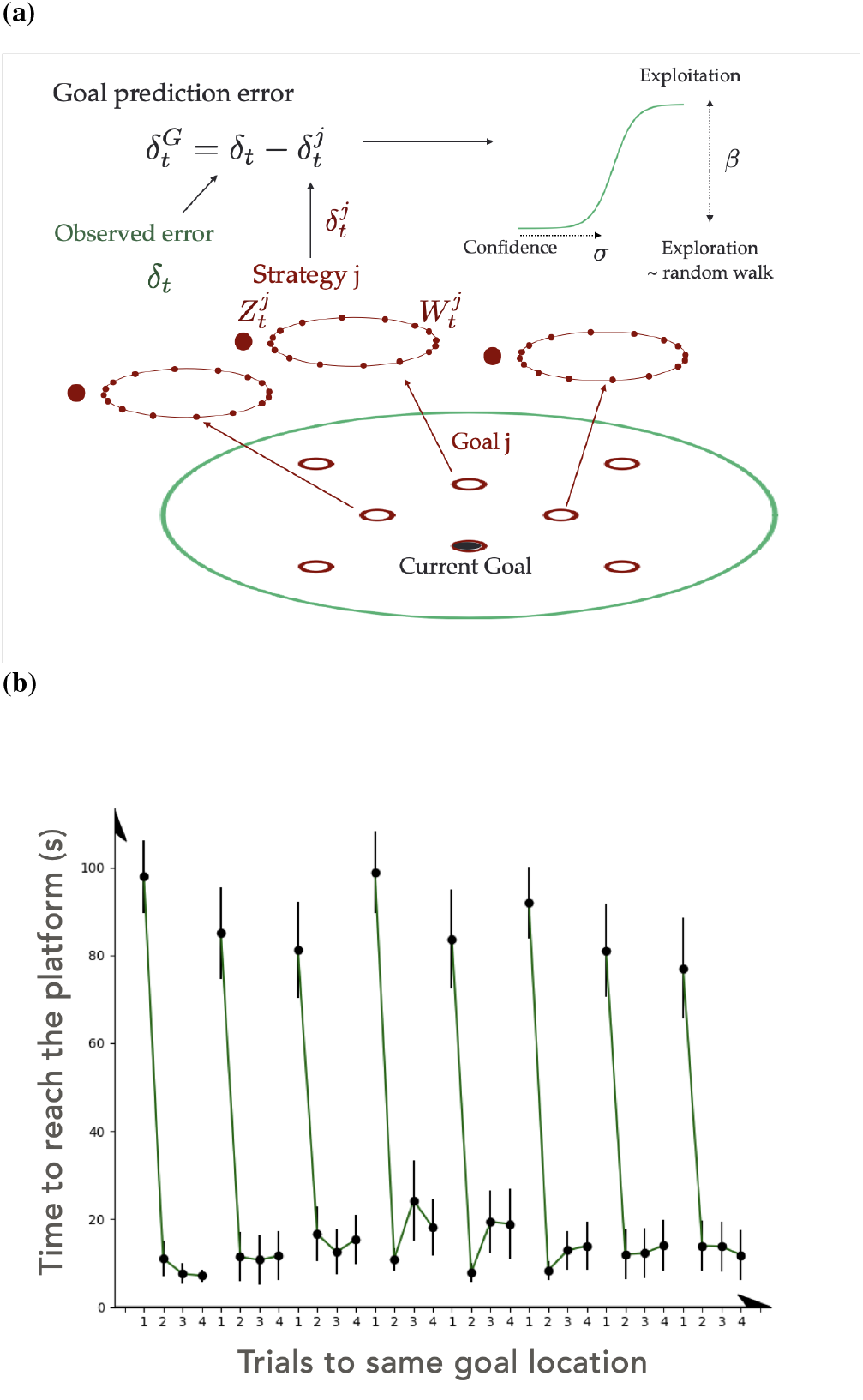
a) Hierarchical RL model. The agent has learned the critic and action connection weights (*Z^j^* and *W^j^*, respectively) for each goal *j* red circles around the maze. The actor and critic networks together, as represented in Fig.3a, form the strategy *j*. A goal prediction error *δ_G_* is used to compute a confidence parameter *σ*, which measures how good the current strategy is in reaching the current goal location. The confidence level shapes the degree of exploitation of the current strategy *β* through a sigmoid function of confidence. When the confidence level is very high, the strategy chosen is closely followed, as shown by a high exploitation parameter *β*. On the contrary, a low confidence level leads to more exploration of the environment. b) Performance of the hierarchical agent. The model is able to adapt to changing goal locations, as seen in the reduction of latencies to reach the goal.

Prefrontal areas have been proposed to carry out meta learning computations, integrating information over multiple trials to perform computations related to a rule or a specific task (Wang et al. 2018). Neurons in prefrontal areas seem to carry goal information (Poucet and Hok 2017;Hok et al. 2005), and their population activity dynamic correlates with the adoption of new behavioural strategies (Maggi et al. 2018). In previous work, prefrontal areas have been modelled as defining the state-action-outcomes contingencies according to the rule requirement (Rusu and Pennartz 2020; Daw et al. 2005). Moreover, prefrontal dopaminergic activity affects flexibility towards changing rules (Goto and Grace 2008; Ellwood et al. 2017), and frontal dopamine concentration increases during reversal learning and rule changes (van der Meulen et al. 2007). Therefore, the goal prediction error that shapes which goal location is pursued according to our hierarchical RL model could be computed by frontal areas from dopaminergic signals.

#### 4.3.1 Limitations in accounting for open field DMP performance

We present this approach as an illustration of how a hierarchical agent could be more flexible by separating the computation of the choice of the goal from the computation of the choice of the actions to reach it.

However, the model has several features that limit its use to provide a neuropsychologically plausible explanation of the computations underlying DMP performance in the watermaze and related open-field environments. First, the agent has to learn beforehand the connections between place cells and action and critic cells that lead to successful navigation towards every possible goal location of the maze. This would involve pretraining with the possible goal locations, whereas the agent would fail to learn a completely new goal location within 1 trial (i.e., return to a location that contained the goal for the very first time). Hence, the model can be considered as a model of one shot recall, rather than one-shot learning. This cannot account for the one-trial place learning performance shown by rats and human participants on DMP tasks towards new goal locations, rather than familiar ones (Bast et al. 2009; Buckley and Bast 2018). One discrepancy between the model’s behaviour and rats can be seen during the first few trials, in which the agent automatically shows adaptation to the new goal location (as reflected by sharp latency reductions from trial 1 to 2; Fig. 5b), whereas rats need a few trials to learn the task (Fig. 1b). However, the hierarchical RL model may account for one-trial place learning performance on the DMP task when the changing goal locations are familiar goal locations, i.e. always chosen from a limited number of locations (Whishaw (1985), see the third point below).

Second, if the agent does not find the goal in the location to which its current strategy leads, it starts exploring the maze randomly until it finds the current goal location and selects a strategy that predicted it the best. This results in consistently high latencies during the first trial of every new goal location (Fig.5b). In contrast, the trial 1 latencies of rats are more variable (Fig.1b), for reasons considered in section 4.2 (last paragraph). On probe trials, removing the goal location would lead the agent to start exploring, therefore failing to reproduce the search preference as shown in open field DMP tasks in the watermaze (Bast et al. 2009), virtual maze (Buckley and Bast 2018) and in a dry-land arena (Bast et al. 2005). Interestingly, this suggests that rats may not show search preference during probe trials when they are tested on a DMP task variant that uses familiar goal location (Whishaw 1985) and, therefore, may be solved by a hierarchical RL mechanism.

Third, a lesion study (Jo et al. 2007), as well as our own inactivation studies (McGarrity et al. 2015), in rats indicate that prefrontal areas are not required for successful one-shot learning of new goal locations, or the expression of such learning, in the watermaze DMP task, and frontal areas were also not among the brain areas where EEG oscillations were associated with virtual DMP performance in our recent study in human participants (Bauer and Bast 2020). This contrasts with the hierarchical RL model, which implicates “meta-control” processes that may be associated with the prefrontal cortex. However, on a DMP task variant that uses familiar goal location (Whishaw 1985) and, therefore, may be solved by this hierarchical agent, prefrontal contributions may become more important, a hypothesis that remains to be tested. This suggests that the two DMP variants may rely on different neuro-behavioural mechanisms. Interestingly, the prefrontal cortex and hippocampo-prefrontal interactions are required for one-trial place learning performance on radial maze (Seamans et al. 1995; Floresco et al. 1997) and T-maze (Spellman et al. 2015) DNMP tasks, which involve daily changing familiar goal locations and, hence, may be supported by hierarchical RL mechanisms similar to our model (see section 4.3). Moreover, on the T-maze DNMP task,Spellman et al. (2015) found that hippocampal projections to mPFC are especially important during encoding of the reward-place association, but less so during retrieval and expression of this association. This is partly in line with the behaviour of the model, as the goal prediction error is important to select the appropriate strategy when the agent finds the correct goal location during the sample trial. However, hippocampal-prefrontal interactions are not yet considered in the model.

Fourth, hippocampal plasticity is required in open field DMP tasks (Steele and Morris 1999;Bast et al. 2005). The current approach suggests that the adaptation necessary during trial 1 gives rise to the selection of a set of actor and critic weights that lead to the goal through the computation of a goal prediction error. The model does not explain how the computation of the goal prediction error would be linked to hippocampal mechanisms. It is possible that a positive prediction error would make the current event (being in the right goal location) salient enough to affect its neural representation to stay in memory, for example. In a spatial navigation task in which rats had to remember reward locations chosen according to different rules,McKenzie et al. (2014) have shown that hippocampal representations are hierarchical depending on the task requirement: if the context was determining the reward location, the context would be the most discriminant factor within hippocampal representations. Recent work Sanders et al. (2020) suggests that hierarchical inference could be used to explain remapping processes. It may be that a hierarchical representation of the task within the hippocampus could help adaptation to new goal locations through remapping processes.Hok et al. (2013) found that prefrontal lesions decreased variability of hippocampal place cell firing and hypothesised that this was linked to flexibility mechanisms and rule-based object associations (Navawongse and Eichenbaum 2013) within hippocampal firing patterns. This finding shows that the prefrontal cortex can modulate hippocampal place cell activity. If the goal prediction error is coded by the prefrontal cortex, these findings imply that the goal prediction error could act on hippocampal representations in order to incorporate new task requirements (*e.g*. information about the new goal location) and modify expectations.

#### 4.3.2 A potential account of arm-maze DNMP performance?

Although the hierarchical RL approach may be limited in accounting for key features of performance on DMP tasks using novel goal locations, it may have more potential in accounting for flexible trial-dependent behaviour displayed by rats on DNMP tasks in the radial arm maze, which involve trial-dependent choices between familiar goal locations. DNMP performance in radial maze tasks requires NMDA receptors, including in the hippocampus, during pretraining, although after pretaining, and contrary to the watermaze (Steele and Morris 1999) and event arena DMP tasks (Bast et al. 2005), rats can acquire and maintain trial-specific place information independent of hippocampal NMDA receptor mediated plasticity, even though the hippocampus is still required (Lee and Kesner 2002; Shapiro and O’Connor 1992; Caramanos and Shapiro 1994). The hierarchical RL architecture may account for this phase of acquisition of arm-reward association via pretraining to the eight possible goal locations, via the formation of actor and critic weights of every strategy. However, the plasticity considered in the hierarchical model is more consistent with changes in hippocampal-striatal connections, whereas the model does not address the role of plasticity within the hippocampus during this phase. Moreover, the hierarchical RL model also fits with the requirement of the prefrontal cortex for flexible spatial behaviour on arm-maze tasks, as described in the previous section.

However, to test if a hierarchical RL architecture can reproduce behaviour on DNMP arm-maze task, our implementation of a hierarchical RL model outlined above (see Fig.5a) would need to be adapted to the arm-maze environments, to the DNMP rule and to an error measure of performance that is typically used in arm maze tasks (Seamans et al. 1995;Floresco et al. 1997). The goal-prediction error would drive exploration to other arms and provide a long term control allowing to carry memories of previously visited goals.

## 5 Conclusions

We presented an actor-critic architecture, which leads to action selection based on the difference of estimated rewards to be received. The model (Foster et al. 2000) uses an estimate of the value function over the maze to drive behaviour, through a critic network that receives place cell activities as input. It can successfully learn to select which action is best through an actor network, which also receives place cell input and is trained to follow the gradient of the value function from the critic difference in successive activities. This agent can follow trajectories towards a particular, fixed, goal location, that corresponds to the maximum of the value function. However, when the goal location changes, the model needs many trials to adjust in order to accurately navigate to the new goal, which is in marked contrast with real DMP performance of rats (Fig.1b) and humans (Buckley and Bast 2018).

To account for one-trial place learning performance on the DMP task, a possible extension to the actor-critic approach is to learn map-like representation of locations throughout the maze that facilitate the direct comparison between the goal location and the agent’s location. This enables the computation of a goal-directed displacement towards any new goal location throughout the maze and reproduces flexibility shown by humans and animals towards new goal location, as reflected by sharp latency reductions from trial 1 to 2 to a new goal location and marked search preference for the new goal location when trial 2 is run as probe (Fig.4b; 4c).

Given that the striatum has been associated with actor-critic mechanisms (Joel et al. 2002;Khamassi and Humphries 2012; O’Doherty et al. 2004;van Der Meer and Redish 2011), using an actor-critic agent for flexible spatial navigation is consistent with empirical evidence associating striatal regions with place learning performance on both incremental (Annett et al. 1989; Devan and White 1999; Braun et al. 2010) and DMP ((Seaton 2019) and Bauer and Bast (2020)) tasks. However, contrasting with the coordinate extension to the actor-critic architecture, experimental evidence suggest that goal location memory may lie within hippocampal place cell representations (McKenzie et al. 2013; Dupret et al. 2010) and that one-trial place learning performance on DMP tasks in rats requires NMDA receptor dependent LTP-like hippocampal synaptic plasticity (Steele and Morris 1999; Bast et al. 2005).

Finally, we illustrated how flexibility may be generated through hierarchical organisation of task control (Balleine et al. 2015; Botvinick et al. 2009; Dayan and Hinton 1993). In an extension of Foster et al. (2000), we separated the selection of the goal from the control of the displacement towards it by means of a different readout of the TD error by the different control systems. The agent first learns different “strategies”, each of which correspond to a different critic-actor component that leads to one of the possible goal locations. The critic and actor are used to perform the displacement to the goal location. An additional hierarchical layer is added to compute a goal prediction error which compares location of the goal predicted from the strategies to the real goal location and selects a strategy accordingly. The agent follows its strategy depending on a confidence parameter that integrates goal prediction errors information over multiple trials. This hierarchical RL agent can adapt to changing goal locations, although these goal locations need to be familiar, whereas in open field DMP tasks the changing goal locations are new (Bast et al. 2005, 2009). However, the hierarchical RL approach may be more suitable to account for situation when trial-specific memories of familiar goal location need to be formed, as on arm-maze DNMP tasks (section 4.3.2)

To conclude, elements of an actor-critic architecture may account for some important aspects of rapid place learning performance in the DMP watermaze task. Together with a map-like representation of location, an actor-critic architecture can support the efficient, fast, goal-directed computations required for such performance, and a hierarchical structure is useful for efficient, distributed, control. Future models of hippocampus-dependent flexible spatial navigation should involve LTP-like plasticity mechanisms and goal location representation within the hippocampus, which have been implicated in trial-specific place memory by substantial empirical evidence.

## Acknowledgements

We thank Georg Raiser for fruitful discussions and comments on the manuscript. C.T. was funded by a PhD studentship supported by the School of Mathematical Sciences and the School of Psychology and the University of Nottingham.

## A Actor-critic equations, based on (Foster et al. 2000)

### Place cell activity

The *N_P_* place cells have Gaussian activation profiles:

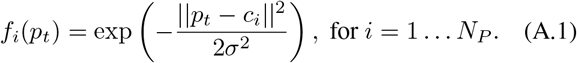

Here 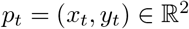 represents the position of the rat at time 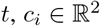 denotes the centre of the ith place cell and *σ* the width of the activity profile.

### Critic

The agent has an estimate of the value of its current state. This estimation is provided by a critic cell, whose activity is computed according to:

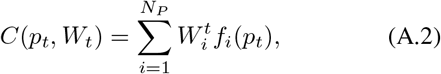

where 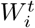 is the weight from the ith place cell to the critic cell.

### Actor

The actor network is composed of *N_A_* = 8 units, which represent 8 possible directions (north, north-east, east, southeast, south, south-west, west and north-west). The activity of the *j*th action cell is computed according to:

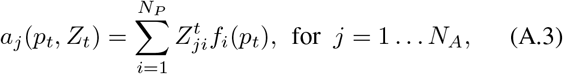

where 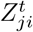 is the connection weight form the ith place cell to the *j* th action cell.

To determine the probability of taking the *j*th action, given the activities *a_j_*, a softmax probability distribution of the activities is computed according to:

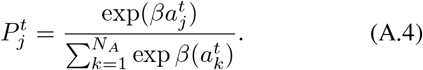

In (A.4), the parameter *β* acts as an inverse temperature. Small values lead to a more uniform probability distribution, increasing randomness, and therefore promoting exploration of the environment. Larger values lead to more exploitation of known state-action pairs.

### Evolution of position

After the selection of the *j*th action, the animal moves according to:

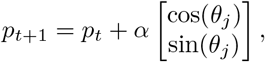

where *θ_j_* denotes the angle of the selected *j*th action cell, and *α* is the speed of the agent.

### Reward

After having moved to the location *p*_*t*+1_, the agent receives a reward *r*_*t* = 1_ if it has reached the goal location (representing the platform in a watermaze experiment), and *r*_*t* = 0_ otherwise.

### TD error

The agent can compare the value of its current state against its prediction through the following TD error:

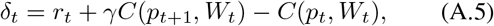

where *C*(*p*_*t*+1_, *W_t_*) refers to the estimated value of the current state *p*_*t*+1_ from the weights at the previous time *t* (see equation (A.2)).

### Learning

The error (A.5) is used to improve the estimator by updating the weights from place cells to the critic (A.2) via 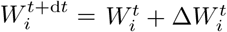, using:

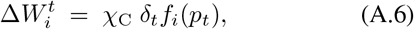

where *χ*_C_is the learning rate. The actor network is also updated in order to improve the policy. The actor weight *Z_ji_* between the *j*th action cell and the *i*th place cell evolves according to 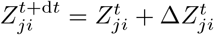, using:

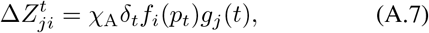

where *χ*_A_is the learning rate, *g_j_*(*t*) = 1 if the *j*th action is chosen and is 0 otherwise. The connection between a place cell and an action cell is strengthened if taking the action at the given location results in an increase in the value function, and is weakened if this results in a decrease in the value function.

### Parameters

To give appropriate feedback to the actor, the critic must be accurate. To help achieve this, it is useful to set the critic learning faster than the actor. This way, the TD error can more rapidly reflect the actual change in value between two consecutive states. Here we choose *χ*_Actor_ = 0.01 and ^χ^_Critic_ = 0.1.

The discount factor *γ* must be quite high to ensure backpropagation of information to the boundaries of the maze. Here we choose *γ* = 0.98.

All the other parameter values are chosen as in Foster et al. (2000).

## B Actor-critic component: effect of changing the place cell activity width and the discount factor on incremental learning towards the goal location

*σ* affects the generalisation between states: if a decision leads to an update in value and policy at a certain state, this decision also updates the value and policy at neighbouring states, the extent to which is weighted by *σ*. In the case where the place cell activity profile is wide, the value function peak is broad (Fig.6a). The agent cannot reach optimal performance, as can be seen from the latencies, that decrease through time (Fig.6b), showing that the agent still is able to learn, but the latencies obtained exceed those for intermediate choices of *σ*. For very narrow place cell activity profiles, the information provided by the value function remains concentrated near the goal location only. The updates cannot be generalised across the space. Therefore it does not provide information of utility for the agent to navigate successfully to the goal from more distant locations.

**Figure 6.**
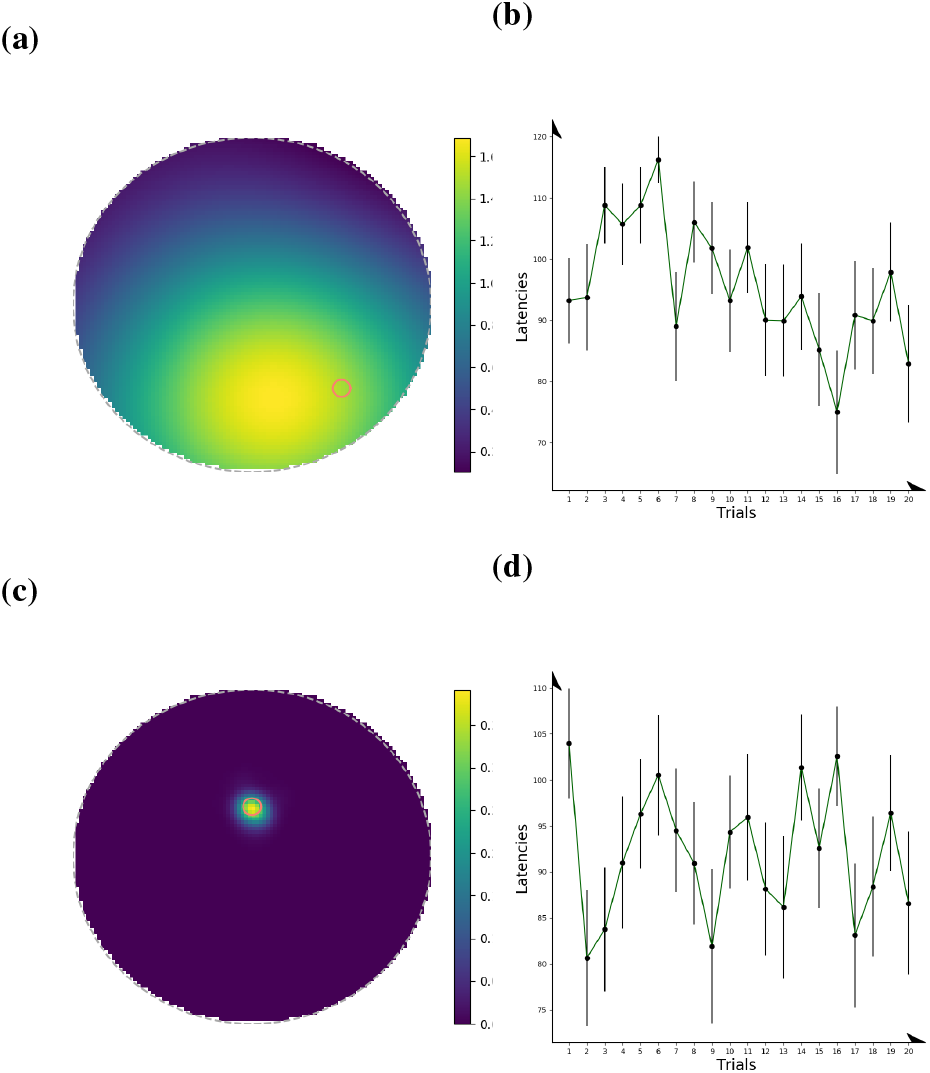
Effect on goal location learning of changing the width of the place cell activity distribution. Value functions (a,c), and latencies (b,d), for high *σ* (a,b) and low *σ* (c,d). A too high value of *σ* leads to a less sharp increase in the value function. The agent learns slightly, but its performance is limited by the opposing updates that it gets when approaching the goal form different directions. A low value of *σ* gives a narrow value update, not allowing any generalisation about which action is optimal. The agent needs many more trajectories to the goal to learn the policy from any starting location.

The discount factor defines how much the estimation of the future value function is taken into account in comparison of the estimation of the current value in the TD error. A low *γ* leads to consequent update only when the agent reaches the goal. The agent is not able to backprogagate the information about the reception of the reward to previous states. This leads to the value function having a steep increase close to the goal location (a), whereas the optimal value function (c) requires a progressive gradient everywhere to guide the actor. The actor therefore does not have a clear information about where to go when it is away form the goal. This is reflected by latencies that stay large (Fig.7b), showing that the agent does not learn the task.

**Figure 7.**
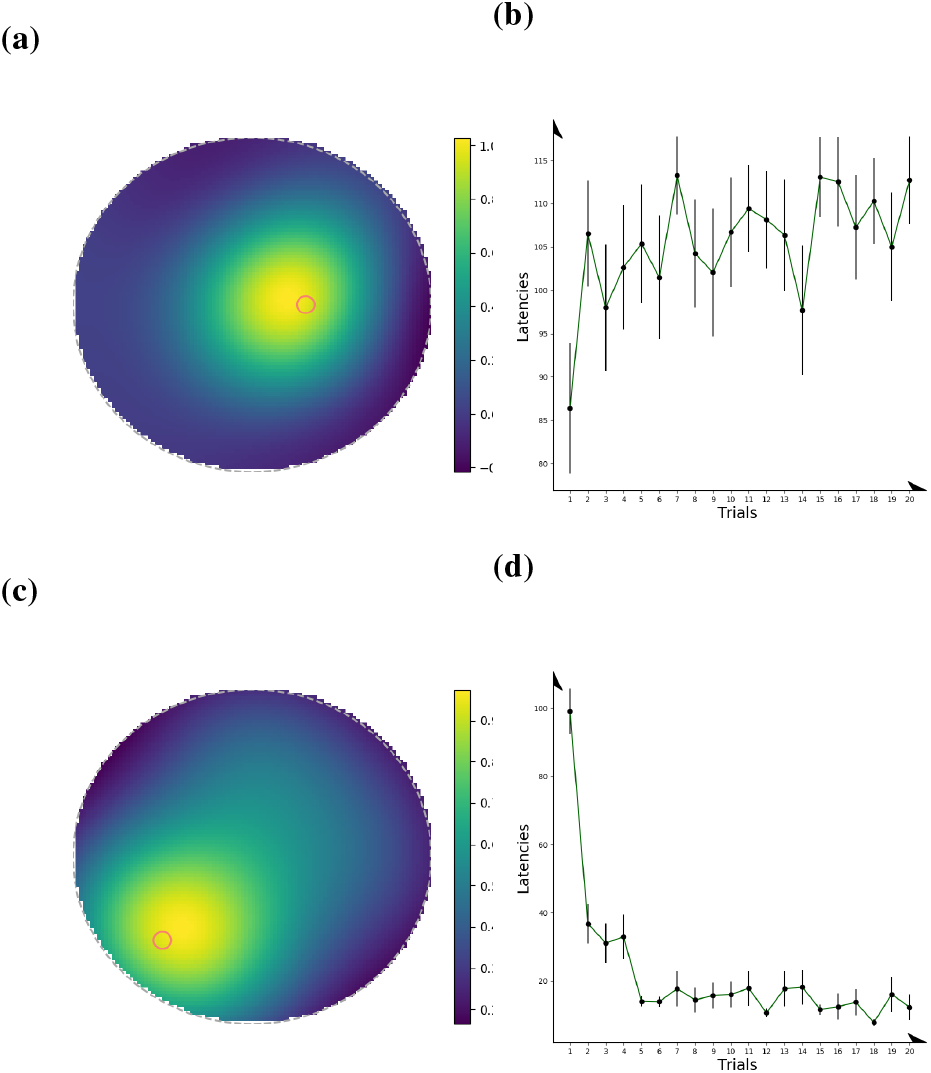
Effect of the discount factor on learning. Low *γ* (a,b) compared to optimal *γ* (c,d), quite high value (0.98). Too low *γ* leads to more narrow value function (a). The agent is not able to backprogagate to previous states about the value of the future state. This leads to the agent not having enough information about optimal actions at the borders of the maze, and prevents learning, as can be seen on the latencies (b). The optimal learning rate in this task should be high enough so that the information is transmitted as fast as possible to the borders of the maze, but cannot be 1 otherwise the same value will be reached across the whole maze, and all actions would be equally optimal.

**Figure 8.**
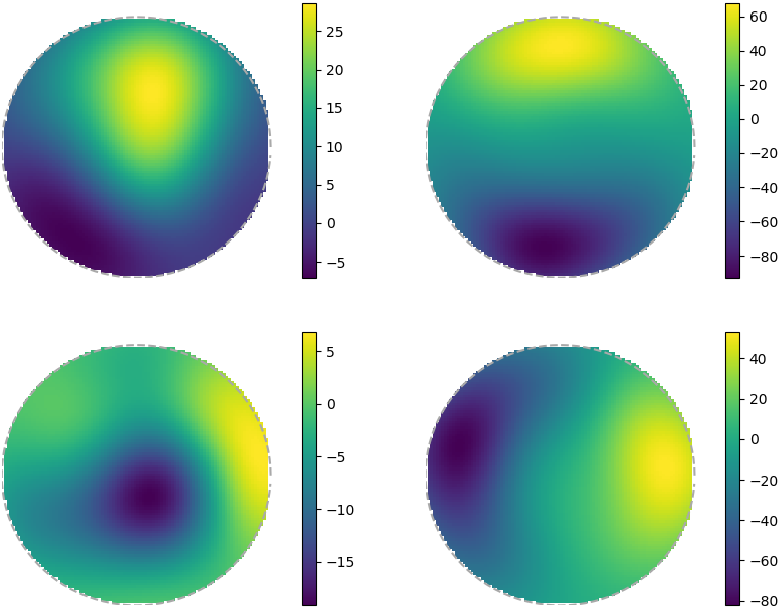
Coordinates estimation throughout the maze. Up: 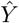, Down: 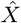. Left: very early on learning, the coordinates are not consistent throughout the maze. Right: after learning, the estimates follow a consistent gradient compared to the real coordinates, which enables correct computation of the motion toward the goal.

### Codes availability

All codes implemented in Julia are available at https://github.com/charlinetess/FMD_SMT

## C Coordinate navigation - equations and supplementary figures

### C.1 Equations

In this section, we describe the equations used to compute the estimates of the *x* and *y* coordinates, as described in section 4.2. Similarly to section A, the positioning system uses place cell activity as input to compute 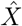 and 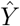, the estimate of the current *x* and *y* coordinates at every timestep:

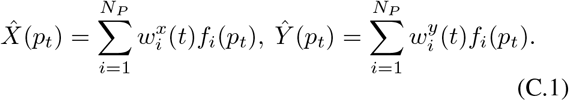

this estimator augments the original model presented in section A. A coordinate action cell computing the direction towards the goal location is added to the existing action cells (see figure 4a). The weights between the position cells and the place cells are updated using self-motion information, according to 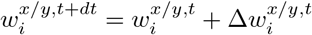, using:

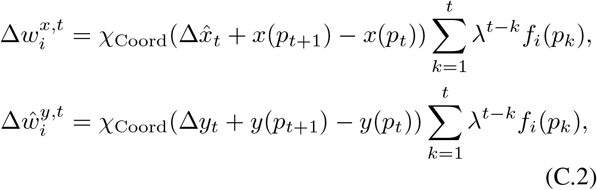

in which *χ*_Coord_ defines the learning rate for the coordinates, and 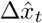 and 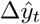 represent the self-motion estimate in the *x* and *y* directions, *i.e*. the difference between the estimated position at the previous location and at the current position, using the previous estimator. They are computed according to:

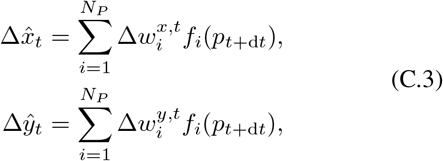

and *x*(*p*_*t*+1_) − *x*(*p_t_*) and *y*(*p*_*t*+1_) − *y*(*p_t_*) refers to the real displacement along the *x* and *y* direction. The term 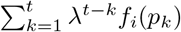 acts as an eligibility trace (Sutton and Barto 2018), adding more importance to the most visited locations. Every time the goal location is encountered, its estimated position 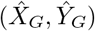 is stored (see figure 4a). Thus, during the subsequent trials, the agent has an estimate of its current location and of the location of the goal at every time step.

To perform navigation, a ninth action cell is added to the original set of eight cells (see figure 4a). Its activity is computed according to equation (A.3) and the probability of selecting it according to equation (A.4). However, contrary to the other direction cells associated to angles that represent different cardinal directions, this action cell is not linked to any particular angle, but drives the agent towards the estimated platform location. If we denote 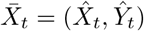 the current estimated coordinates and 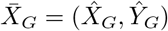 the current estimate of the goal location, the selection of the coordinate action induces the following change in position:

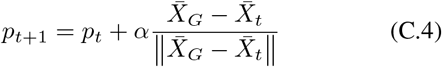

A further distinguishing feature of this action cell is in the update of weights linked to this cell. The learning rule does not depend on position; instead, the weight between the 9th action cell and the *i* place cell is computed according to 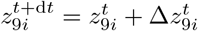:

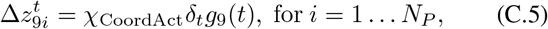

where *g*_9_(*t*) = 1 if the 9th action cell has been selected and zero otherwise, and *δ_t_* refers to the error computed from equationA.5. When there is no goal coordinate in memory, the direction is chosen at random among the 8 other directions and the coordinate action cell weights are not updated. All of the other weights *z_ji_, j* = 1…8, *i* = 1…*N_P_*, evolve according to equation A.7.

#### Parameters

Discount factor: = 0.99, Learning rate for coordinates : *χ*_Coord_ = 0.05, Eligibility trace parameter: λ = 0.8.

#### Codes availability

All codes are implemente din Julia and available at https://github.com/charlinetess/FMD_DMP

### C.2 Supplementary figures

## D Hierarchical model - equations

In this approach, the agent learns the weights corresponding to 8 different goal positions throughout the maze, using the model from Appendix A. This gives rise to 8 possible strategies, each providing a control mechanism to produce trajectories to a given location.

At the start, a strategy *j* is selected at random. The observed error *δ_t_* is the error computed from the critic of the *j*-th strategy but the reward component refers to the real reward received, whereas the error computed from the strategy *j* refers to the predicted reward. Symbolically, we have that:

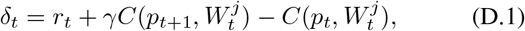

and

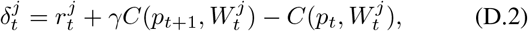

where 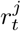 refers to the predicted rewards according to strategy *j*, and *r_t_* refers to the reward available in the environment. The strategy prediction error if currently following strategy *j* is computed from the difference of the two errors:

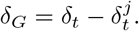

The strategy prediction error is used to shape the dynamic of a confidence level *σ_t_* that evolves according to:

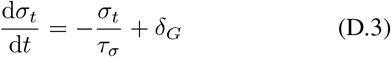

The temporal evolution captures the fact that error in predictions should lead to a transient reduction in the confidence of the agent in this particular strategy. The confidence level determines how the agent’s actual behaviour respects the defined policy. The temperature parameter from equation (A.4) now becomes dependent on the confidence level:

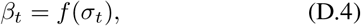

where *f* is a Sigmoid function given by:

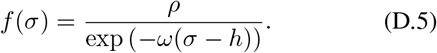

Here, *ρ* defines the gain factor, *h* the cut threshold (point of highest slope in the sigmoid). *ω* defines the steepness of the function around the cut threshold, which shapes how fast the exploitation parameter evolves when the confidence parameter becomes higher or lower than the cut threshold. When the goal is not found in the location estimated by the followed strategy, the confidence parameter decreases, allowing exploration of the environment, until the goal is eventually found. When the goal is found, the strategy *j* which minimises the goal prediction error is selected. This leads to the confidence parameter to start ramping up again.

### Parameters

We use *ω*=5, *h*=-0.2, *ρ*=2, *τ_σ_*=15.

### Codes availability

All codes implemented in Julia are available at https://github.com/charlinetess/MetaTD

